# Quantitative Expansion Microscopy for *In Situ* Estimation of Endogenous Target Abundance

**DOI:** 10.64898/2026.01.18.700178

**Authors:** Matthew D. Lycas, Juan C. Landoni, Benedetta Noferi, Christian E. Zimmerli, Kyle M. Douglass, Suliana Manley

## Abstract

Spatially mapping protein abundance *in situ* can offer important insights into molecular mechanisms and the physiological functions of protein complexes. This is typically achieved by combining super-resolution microscopy to image fluorescently-labeled protein locations with statistical estimators to retrieve abundances, where accuracy is strongly impacted by labeling efficiency. We introduce quantitative expansion microscopy (qExM) as a method to estimate endogenous protein abundance on ExM data, which offers improved antibody targeting through molecular decrowding. Using cryo-fixation, we preserved ultrastructure and enhanced labeling efficiency to improve accuracy in abundance estimations. We benchmark the effectiveness of qExM by quantifying the stoichiometry of well-characterized nuclear pore complex subunits, and find a mean percent error of 9.4%. We further apply qExM to investigate the abundance of mitochondrial respiratory chain complexes in functionally distinct organelle subpopulations, and of mitochondrial respiratory chain super-complexes in differentially activated human T-cells. qExM provides a robust methodological framework for quantifying endogenous protein abundance in expanded samples *in situ*.

## Introduction

Historically, molecular biology has benefited from bulk biochemical detection to investigate the function and abundance of protein complexes, although lacking valuable spatial information. More recently, super-resolution microscopy unlocked the ability to specifically visualize and resolve protein-scale organization *in* situ, imaging individual complexes and using statistical methods to infer protein abundance^1,2^. Quantitative super-resolution imaging has revealed relationships between neuronal synapse active zone proteins and neurotransmitter release, between CD19 receptor protein expression and myeloma cell targeting by CAR-T immunotherapy, and between therapeutic treatment and CD20 receptor protein organization^3–6^. Such methods are complementary to state of the art electron microscopy techniques like in situ cryo-electron tomography, which can identify molecular complexes via their structures in subcellular regions^7^. To map protein abundances across whole cells, fluorescence super-resolution localization microscopy (SMLM), including PAINT, STORM, and PALM, have been used to resolve individual labeled targets^1,2,8^. Typically, such methods use dual-color images to first estimate labeling efficiency, which is then used to account for underlabeling and infer the number of proteins. Calibration with exogenous constructs of known stoichiometry are typically added to the biological sample to determine labeling efficiency ^1,2,9^, a limiting factor in abundance accuracy.

Expansion microscopy (ExM) has emerged as a unique super-resolution method that physically enlarges specimens while preserving their relative protein organization^10–12^. In addition to increasing resolution, ExM enhances antibody accessibility to epitopes through molecular decrowding, and reduces the effective size of labels relative to the specimen^13–15^. This circumvents issues associated with traditional immunolabeling approaches such as steric hindrance due to target crowding or epitope inaccessibility^16^. Furthermore, since it is a sample-based approach, ExM resolution is improved multiplicatively when combined with optical approaches such as SIM, STED, or SMLM^17–21^. ExM also reduces scattering from thick samples, and thus has been useful for imaging medically relevant biological samples, such as pathological tissue and parasites within hosts, providing super-resolution where SMLM would not be effective^22,23^. However, ExM has thus far not been developed as a quantitative method to estimate protein abundances.

Here, we introduce quantitative expansion microscopy (qExM), which combines optimized ExM protocols with stimulated emission depletion (STED) microscopy and statistical models to estimate abundances of *in situ* target proteins^1,8,18,24,25^. By relying on cryo-rather than chemical-fixation, as well as anchoring with glycidyl methacrylate (GMA), we improve antibody labeling efficiency^24–26^. Then, using ExM-STED images of protein targets labeled with two independent antibodies, we apply statistical models to simultaneously estimate both labeling efficiency and target abundance^1,2,8,27–29^. We benchmark qExM using the well-characterized eight-fold symmetry of the nuclear pore complex (NPC)^30^. We then employ qExM to investigate differences in respiratory chain super-complex abundances between functionally distinct mitochondrial sub-populations, and between non-activated and activated T-cells, offering spatial information absent from bulk biochemical approaches. qExM transforms standard, commercially available antibodies into quantitative tools, broadening the range of target proteins. This approach allows endogenous protein abundance to be mapped *in situ* without genetic manipulations, and lends accessibility to cellular, tissue-level, or organismal contexts.

## Results

### Labeling Efficiency and Abundance Estimates through Quantitative Expansion Microscopy

Accuracy in counting target proteins or protein complexes in fluorescence imaging is most strongly determined by two parameters: spatial resolution and labeling efficiency. Resolution allows targets to be spatially separated, since overlapping targets will be considered as a single object. Labeling efficiency determines the fraction of targets that can be detected by fluorescence. Improvements in super-resolution imaging techniques have resulted in localization precisions on the order of or below single proteins, but low labeling efficiency strongly degrades accuracy and must be accounted for to estimate endogenous target abundances.

An analogous problem in population ecology known as capture-recapture aims to estimate population sizes based on random, incomplete sampling^8^. In these approaches, two subsets of individuals within a population are captured and marked sequentially, and a statistical framework is used to estimate the total population from the sample sizes and the number of recaptured individuals

In a multicolor fluorescence microscopy experiment, spectrally distinct antibody labels can be considered equivalent to the different markings in time, and antibodies against different epitopes or proteins within a complex ensure independent sampling (Fig. 1a). In a two-color experiment there are four possible outcome classes for each target: only label A, only label B, both labels, or no labels; images are segmented and outcome classes identified (Fig. 1b). While our measurement cannot detect the ‘no labels’ class, we can derive estimates of the two labeling efficiencies and the total target abundance (Fig. 1c). Labeling efficiencies are estimated using maximum likelihood estimation (MLE) restricted to only the observable outcomes. A variety of estimators exist for target abundance; previous fluorescence microscopy studies have estimated labeling efficiency in a similar manner, typically using the well-known Petersen estimator^1,2,29,31^. In contrast, we use the Chapman estimator because, by assuming a hypergeometric distribution for the distribution of targets having both labels, it has less bias and avoids numerical errors due to division by zero (Fig. S1, S2)^32^.

**Figure 1.**
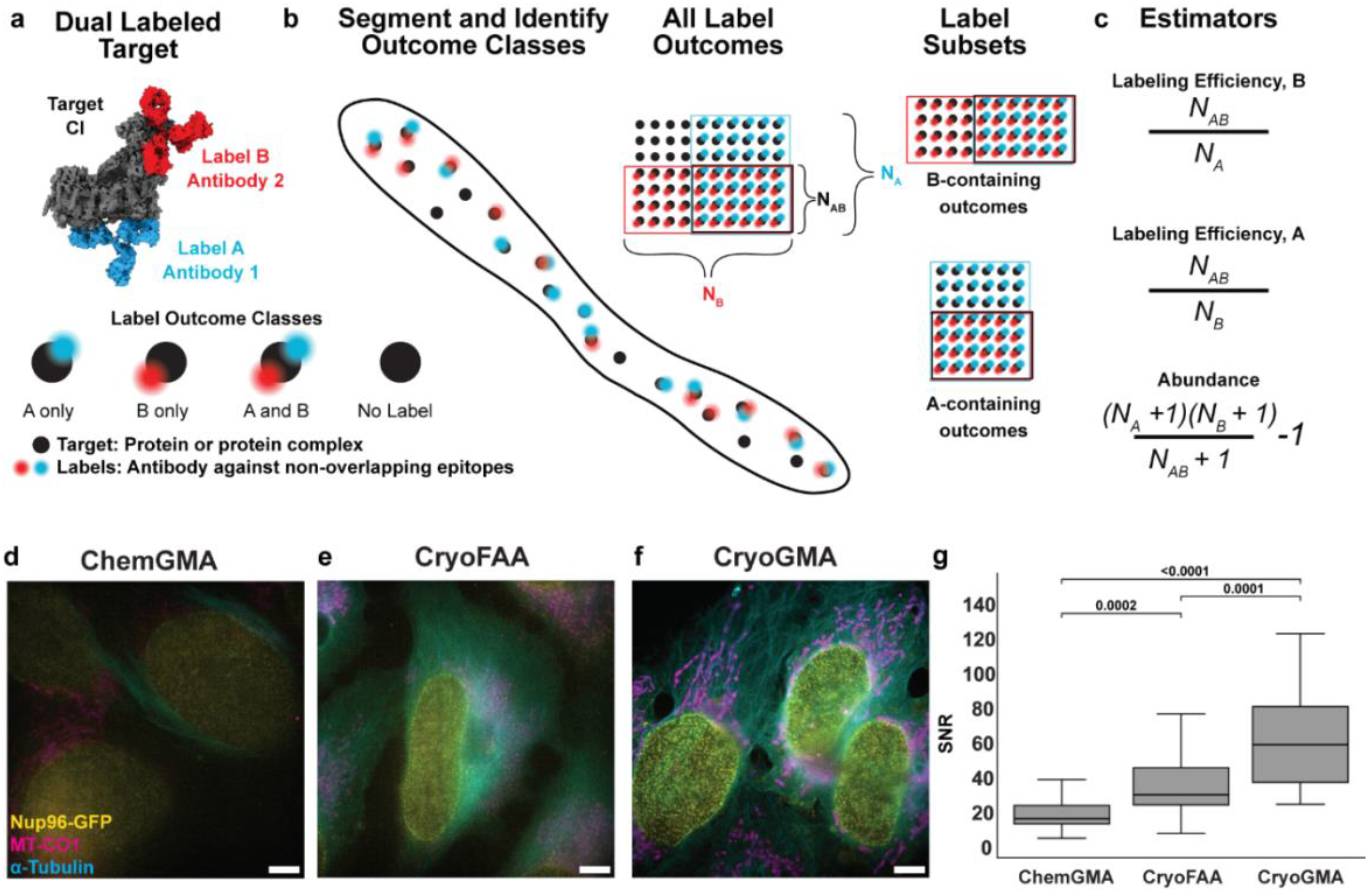
Quantification principles of the capture-recapture approach in qExM. **(a)** The qExM framework targets endogenous proteins or protein complexes using pairs of antibodies against non-overlapping epitopes, leveraging western blot-effective antibodies enabled by expansion microscopy’s improved labeling efficiency. **(b)** Schematic representation of the labeling outcome categories in a qExM experiment. The total population of targets can be labeled by either antibody A (blue), antibody B (red), both antibodies (dual-labeled), or remain unlabeled. Each label provides a random subset of the total population, allowing assessment of the other label’s efficiency. **(c)** Mathematical formulation of the estimators used in qExM to calculate labeling efficiency of each antibody and the total abundance of target proteins, derived from the Chapman capture-recapture method. **(d**,**e**,**f)** Fluorescence microscopy images of U2OS cells prepared using three expansion microscopy protocols and collected and displayed with identical parameters: d: chemical fixation with glycidyl methacrylate anchoring (ChemGMA), e: cryo-fixation with formaldehyde/acrylamide anchoring (CryoFAA), and f: cryo-fixation with glycidyl methacrylate anchoring (CryoGMA). Cells were immunolabeled for Nup96-GFP (cyan), mitochondrial MT-CO1 (magenta), and α-tubulin (yellow). Scale bar 5 µm. **(g)** Signal to Noise measurements comparing the mitochondrial MT-CO1 signal across the three expansion microscopy protocols.

A high labeling efficiency is crucial to ensure a high accuracy of the estimators (Fig. S1). To improve antibody labelling efficiency for qExM, we combined cryo-fixation with GMA anchoring (Fig. 1d,e,f)^24,25^, resulting in an approximately 2-fold increase in the labeling efficiency over protocols employing formaldehyde during either fixation or anchoring (Fig. 1g). This is likely due to improved epitope availability for immunolabelling by avoiding fixation-induced blockage^26^. We also observed a decrease in non-specific antibody labeling on expanded samples compared to non-expanded samples, which will further improve the accuracy that can be expected with qExM (Fig S3).

Increasing the expansion factor offers the possibility to work with targets of higher endogenous density, with smaller distances between targets. We investigated this possibility by reducing the concentration of bis-acrylamide in the ExM monomer mixture^33,34^. However, we needed to employ more destructive denaturing protocols to achieve greater expansion factors, which resulted in a decrease in antibody labeling efficiency^33^ (Fig. S3). We thus settled on an expansion factor of 4.3x as a suitable compromise between antigenicity and spatial resolution (Fig S3, S5).

Finally, as with any image-based counting method, qExM is limited to objects which can be spatially resolved, and we refer to our estimates as “target abundance.” Proteins may exist in multimeric complexes which are not spatially separated; applied to such cases qExM serves to estimate the number of these multimeric complexes. We performed qExM to achieve the following results by combining ExM with STED, bringing our spatial resolution down to ~10 nm^18^.

### Validation of qExM Using Nuclear Pore Complex Octameric Symmetry

We benchmarked qExM against the well-characterized structure of the nuclear pore complex (NPC), whose subunits are arranged with 8-fold radial symmetry. We used CRISPR knock-in NUP96-mEGFP U2OS cells, which offered the advantage of high-efficiency targeting by GFP-antibodies. We implemented three parallel imaging experiments of NPC subunits by targeting NUP96-mEGFP via an antibody against GFP (label A) in combination with a second antibody (label B) against either NUP96, NUP54, or ELYS (Fig. 2a-c). These proteins are known components of the NPC’s octameric arrangement (Fig. 2d), providing a ground truth for our measurements^30,35^.

**Figure 2.**
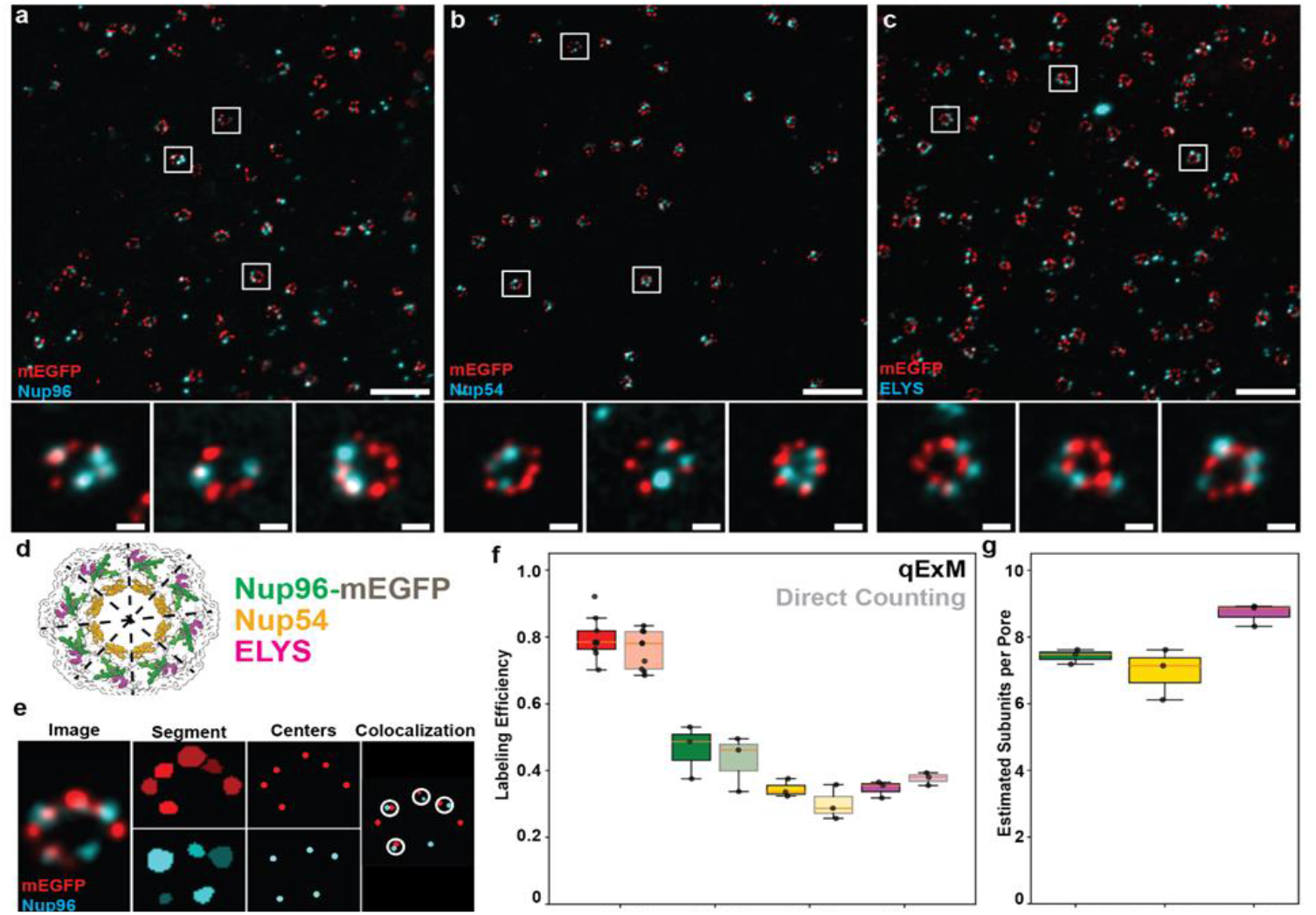
Validation of qExM using the known 8-fold symmetry of nuclear pore complexes. **(a)** qExM STED images of Nup96-mEGFP U2OS cells with mEGFP (red) and Nup96 (blue) labeling. **(b)** qExM STED images of Nup96-mEGFP U2OS cells with mEGFP (red) and Nup54 (blue) labeling. **(c)** qExM STED images of Nup96-mEGFP U2OS cells with mEGFP (red) and ELYS (blue) labeling. Smaller images in a-c show individual nuclear pores at higher magnification, demonstrating the characteristic ring-like arrangement of NPC subunits. **(d)** Structural model of the human nuclear pore complex highlighting the positions of Nup96-mEGFP (green), Nup54 (yellow), and ELYS (magenta) within the 8-fold symmetrical arrangement. **(e)** Image analysis workflow showing raw data (Image), segmentation mask (Segmentation), instance identification (Centers), and colocalization analysis (Colocalization). **(f)** Labeling efficiency estimates from qExM mEGFP and each target protein. “Direct Counting” refers to the labeling efficiency defined by the number of observed target per nuclear pore, knowing that there should be 8 total targets. No statistical significance was found between groups (mEGFP p = 0.23, Nup96 p= 0.65, Nup54 p= 0.28, ELYS p = 0.19) **(g)** Estimated abundance of protein subunits for each experiment, showing means of 7.4, 7.0, and 8.7 for Nup96, Nup54, and ELYS, respectively, closely matching the expected 8-fold symmetry. Scale bars: 500 nm (overview images) and 50 nm (insets). 3 experimental replicates were made for each imaging pair. 45 images of 1365 NPC wlabeled ith Nup96-mEGFP with Nup96. 41 images of 1465 NPC labeled with Nup96-mEGFP with Nup54. 32 images of 1118 NPC labeled with Nup96-mEGFP with ELYS.

We developed an analysis pipeline (Methods) to segment individual NPC subunits in each channel and quantify their colocalization (Fig. 2e)^36^; this allowed us to retrieve the experimentally-determined variables required to apply the Petersen or Chapman estimators. Fluorescence signal was associated with a specific NPC based on its proximity (<82 nm) to its manually-defined center. To determine colocalization between label A and label B, we set distance thresholds between subunits in each channel in an approach similar to that used in qPAINT (Methods) ^1,31^. This data-driven approach to setting the threshold ensured that our colocalization criteria accounted for the experimental labeling conditions specific to each experiment.

We estimated the abundance of NPC subunits by applying the framework without using the labeling efficiency. The number of each outcome class was determined for each image and directly used to compute an abundance estimate (Fig. 1). We then calculated the number of subunits per NPC by dividing the abundance by the number of NPCs in individual images. Considering each labeling condition individually, we obtained abundance estimates of 7.4 ± 0.22, 7.0 ± 0.78, 8.7 ± 0.34 per NPC (errors are the standard deviation from three biological replicates (Fig. 2g)) for NUP96, NUP54, and ELYS, respectively. Thus, percent errors from the expected value of 8 (100 x (abundance estimate – expected value)/expected value) were 7.5%, 12.5%, and 8.75%. The standard deviation of the percent errors decreased from 31.3% to 8.4% when estimation was performed on a whole image instead of on a single NPC. This reflects the power of accumulating a larger number of identified targets to obtain more precise abundance estimates (Fig. S2). Averaging all three labeling targets, the overall abundance estimate is 7.7 ± 0.87, within error of the expected value of 8.

For each pair of labels, we then quantified the number of targets bound by each probe, as well as their colocalization, and used those values to estimate the labeling efficiency of both probes (Fig. 2f). Labeling efficiency in this context is defined as the probability for each subunit of the NPC to be labeled. We further calculated effective labeling efficiencies per protein using the known number of protein copies per NPC (NUP96 = 32, NUP54 = 32, and ELYS = 16), equivalent to the number of binding sites available for antibody binding, as prior information (Fig. S4). These lower labeling efficiencies reflect the reduced likelihood that the label will bind to a single protein. To benchmark the estimated labeling efficiency, we independently measured it by dividing the number of identified targets per NPC by 8; this we refer to as “direct counting” where a value of 1 would correspond to all 8 symmetric units being captured by an image. We found the mean values from direct counting to be statistically indistinguishable from our qExM estimates (Fig. 2f, “Direct Counting”). Thus, our framework offers a highly accurate method for estimating labeling efficiency.

### Quantifying Mitochondrial Respiratory Chain Complex Abundance

Respiratory chain complexes are multi-protein assemblies that reside on the inner mitochondrial membrane, which are responsible for creating the proton gradient that drives oxidative phosphorylation and a wide variety of metabolic pathways. Their abundances are biophysically important since they are a determining factor in ATP production efficiency ^37–39^. Each complex is composed of multiple distinct subunits and assembly factors originating from both mitochondrial and nuclear genomes; thus, the use of genetically-encoded labels presents a risk of perturbing their assembly through disrupted targeting or interference with protein-protein interactions ^40^. To avoid this, we applied qExM with two-color antibody labeling in U2OS cells to the following pairs of proteins: MT-ND2/NDUFS2 for Complex I, UQCRCP1/UQCRFS1 for Complex III, and MT-CO2/MT-CO1 for Complex IV (Fig. 3a-c) ^41^.

**Figure 3.**
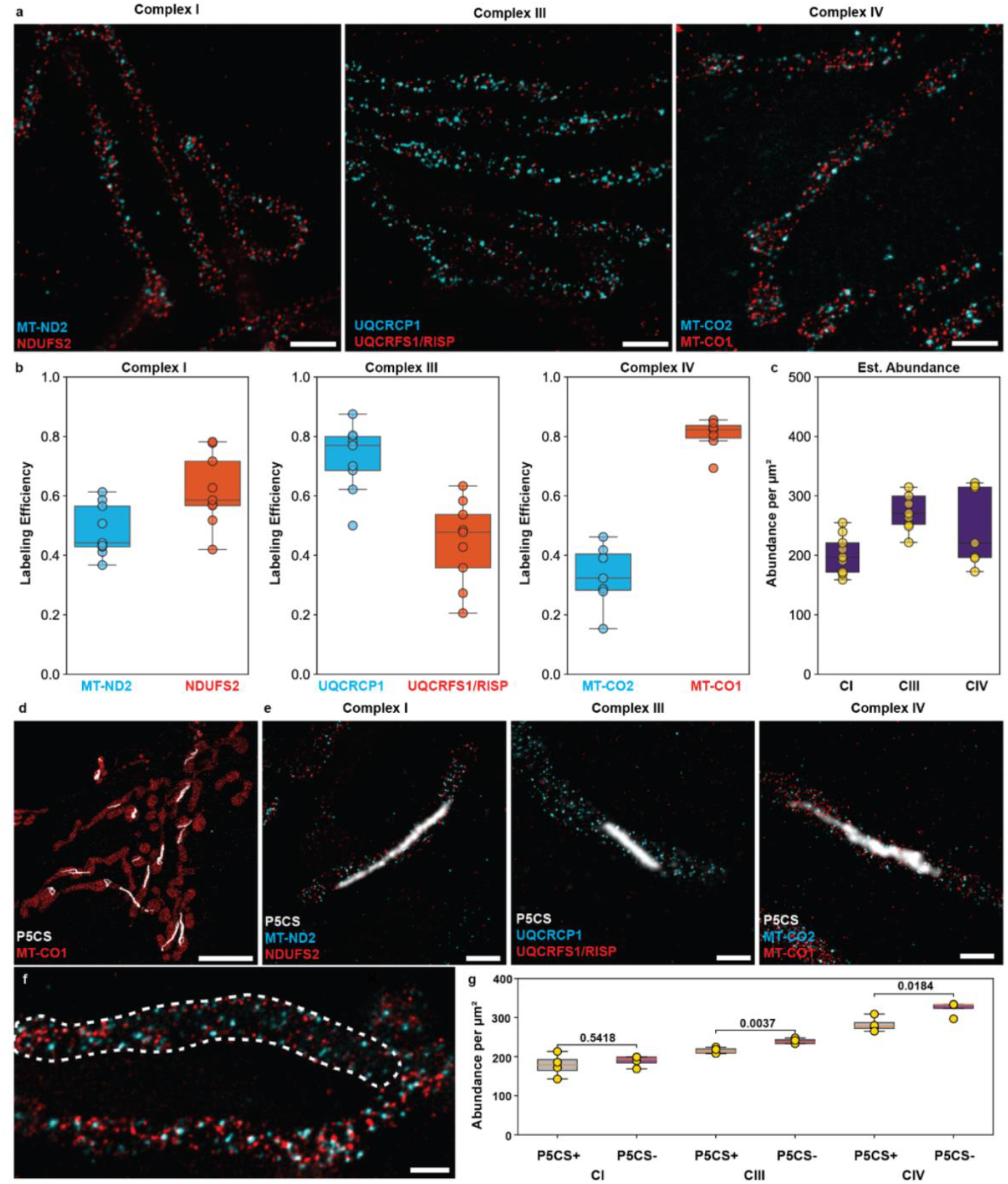
qExM quantification of mitochondrial respiratory chain complexes. **(a)** qExM STED image of respiratory chain Complex I in U2OS cells, labeled for MT-ND2 (blue) and NDUFS2 (red), Complex III, labeled for UQCRCP1 (blue) and UQCRFS1/RISP (red), and complex IV, labeled for MT-CO2 (blue) and MT-CO1 (red). Scale bars: 500 nm. **(b)** Labeling efficiency distributions for each antibody used to target respiratory chain complexes, showing varied efficiencies that highlight the importance of calibration for accurate quantification. **(c)** Estimated abundances per μm^2^ of mitochondrial area for respiratory chain Complexes I, III, and IV in U2OS cells, derived from qExM measurements. 9 experimental replicates (in gold) comprised from 170 images of CI, 114 images of CIII, and 152 images of CIV. **(d)** Dual-color expansion NSPARC image of a nutrient starved U2OS cell labeled for MT-CO1 (red) and P5CS (white). Scale bar: 4 µm. **(e)** qExM images for Complex I, Complex III, and Complex IV with P5CS labeled (white). Scale bar 500 nm. **(f)** Example qExM image for Complex IV ide tifying a mitochondrion Previous estimates of complex densities inSTED n that contained the P5CS fiber (while dashed line). **(g)** Estimated abundances per μm^2^ of mitochondrial area for respiratory chain Complexes I, III, and IV in U2OS cells in and out of P5CS signal, derived from qExM measurements. Data is comprised of 4 experimental replicates with 45 images for CI, 45 images for CIII, 48 images for CIV.

We adapted the analysis methods developed for the NPC as follows to identify individual respiratory chain complexes in dual-color images and determine signal colocalization within mitochondria. Mitochondria were segmented from the Gaussian-blurred summed raw image from both channels by a random forest classifier trained in LABKIT^42^. Then, puncta corresponding to respiratory chain complexes within these masks were found through instance segmentation with CellProfiler^36^, and colocalization thresholds for nearest neighbor distances were calculated following the same procedure described for the NPC.

Following instance segmentation and colocalization analysis, we estimated the labeling efficiency for each antibody (Fig. 3d,e). Antibodies targeting NDUFS2, UQCRCP1, and MT-CO1 showed higher mean labeling efficiencies (0.65, 0.72, 0.80) compared to those for MT-ND2, UQCRFS1/RISP, and MT-CO2 (0.51, 0.47, 0.30), highlighting the importance of antibody selection for quantitative studies. We used the Chapman estimator to estimate the abundance of each respiratory chain complex per unit projected area of mitochondria. In this way, we could compare abundances in normal conditions (Fig. 3e) and under starvation, where mitochondria have been recently reported to form metabolically distinct populations by sequestering pyrroline-5-carboxylate synthase (P5CS)^43^ (Fig. 3f-j). The basal abundance of respiratory chain complexes was similar to that expected from biochemical studies, and respiratory chain complexes III and IV were found to be lower in mitochondria containing P5CS filaments, consistent with the published model of P5CS mitochondria requiring membrane potential but incapable of complete oxidative phosphorylation. These findings highlight the interest in quantifying the abundance of proteins *in situ* with spatial context (Fig. 3j).

Since the size of a respiratory chain complex (~15 nm) ^44^ is slightly larger than our resolution limit (10 nm), we expect abundances estimated by qExM to reflect the densities of complexes. heart mitochondria (413 CI, 572 CIII, 2030 CIV per µm^2^ of cristae) model the number per 2D area of crista membrane^45^. We measured (188 CI, 253 CIII, 261 CIV per µm^2^ projected onto the 2D imaging plane) for mitochondria in U2OS cells, which includes the matrix; thus we expect that this underestimates the density per crista membrane region.

We also would expect a lower density of complexes in U2OS compared with cardiomyocytes due to differences in bioenergetic demand. Previous quantifications of respiratory chain complex density either reported relative changes across experimental conditions or the ratio of the different complexes ^46–48^ or were indirect^45^; in contrast, qExM can offer absolute abundances *in situ*.

### Analysis of Respiratory Chain Super-Complex Formation

Respiratory chain complexes can assemble into higher order super-complexes, an organized structure proposed to improve their efficiency and/or stability^49,50^. Current knowledge of respiratory chain super-complex formation derives primarily from bulk biochemical measurements such as blue native gel electrophoresis and structural techniques including cryo-electron microscopy^44,51,52^, so the composition and abundance variations *in situ* across cellular scales has remained inaccessible. Such information would lend insight into their assembly, spatial heterogeneity, and plasticity across cellular conditions.

To investigate this, we applied qExM to U2OS cells. We performed three-color expansion STED imaging to simultaneously visualize Complex I (NDUFS2), Complex III (UQCRCP1), and Complex IV (MT-CO1), the key components of super-complexes (Fig. 4a,b). We repeated the previous colocalization analysis, with seven possible outcome classes: one complex labeled (CI, CIII, or CIV), two complexes labeled (CI and CIII, CI and CIV, or CIII and CIV), or all three labeled (CI, CIII and CIV). Using these experimental values and the previously determined labeling efficiencies for our protein/antibody pairs, we solved the seven linear equations to estimate the abundance of each class (Fig. 4c).

**Figure 4.**
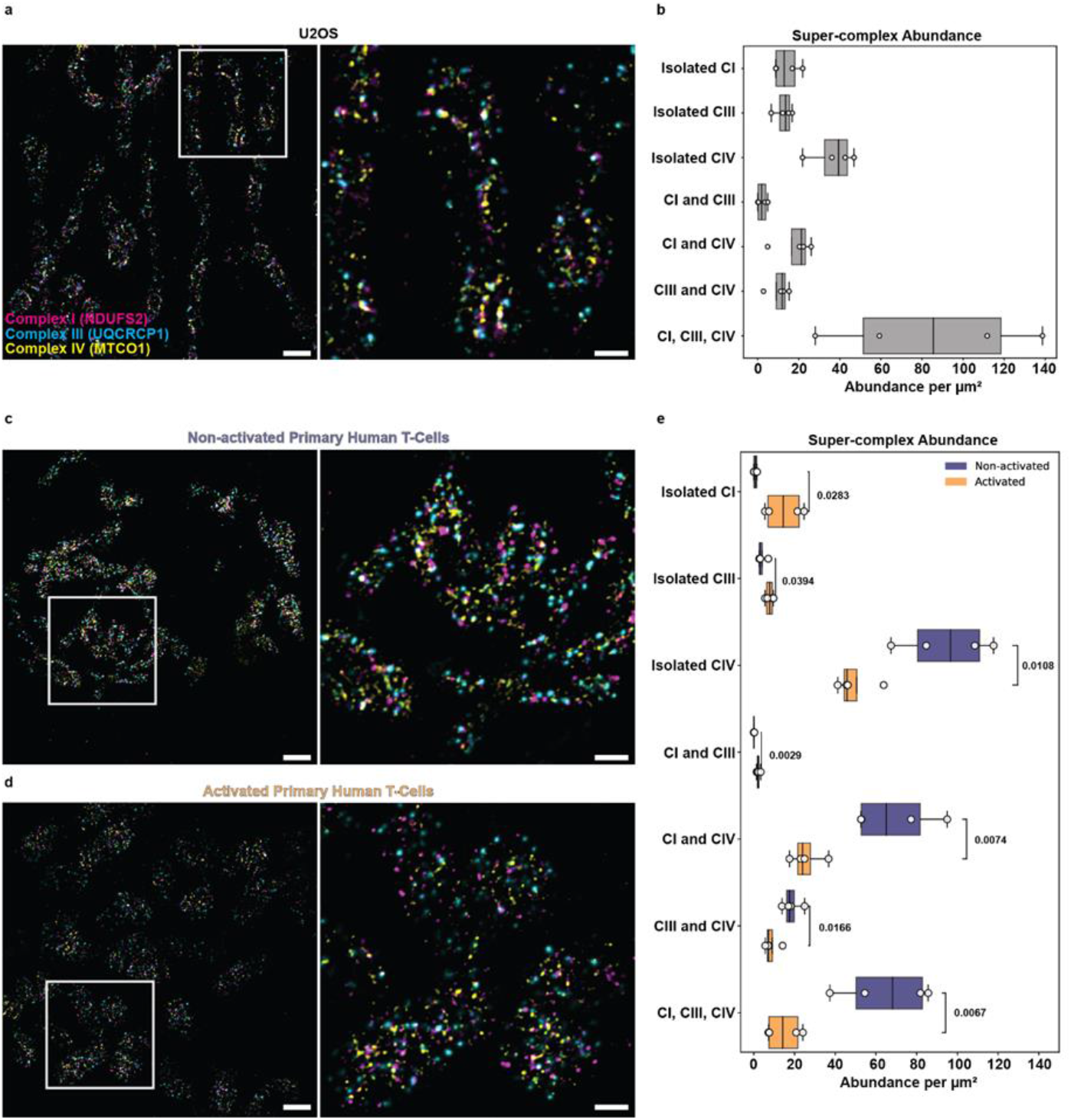
Analysis of respiratory chain super-complex composition in U2OS cells and human T-cells. **(a)** qExM STED image of U2OS mitochondria labeled for Complex I (NDUFS2, magenta), Complex III (UQCRCP1, blue), and Complex IV (MT-CO1, yellow). **(b)** Quantification of labeling efficiency-corrected abundances for isolated complexes and various super-complex configurations in U2OS cells, revealing the predominance of CI+CIII+CIV assemblies. Data originates from 4 replicates containing 34 images total. **(c)** qExM STED image of non-activated primary human T-cells labeled for respiratory chain components. **(d)** qExM STED image of activated primary human T-cells showing altered distribution of respiratory complexes. **(e)** Comparison of abundances of isolated complexes and super-complex configurations between non-activated (purple) and activated (orange) T-cells, showing significant reduction in super-complex formation upon activation. Scale bars: 500 nm (left a, left c e, left d) and 200 nm (right a, right c, right d). Data originates from 3 independent patient samples made into 4 total replicates, 40 images of activated T-cells and 44 images of non-activated T-cells.

Consistent with previous biochemical studies^51,53^, our analysis revealed a predominance of Complex I+III+IV super-complexes in U2OS cells, with most of Complex 1 and Complex 3 assembled into super-complexes, and Complex 4 both isolated and in super-complexes^51^. In contrast, we observe a decreased proportion of individual Complex 4 relative to super-complexes compared with published values in U2OS cells^51^. We speculate that detergent-based protein isolation could partially disassemble super-complexes during biochemical preparations. Intriguingly, we identified spatial heterogeneity in super-complex density, with globular regions of ~500 nm in diameter containing a high concentration of super-complexes (Fig. S6).

During T-cell activation, cells undergo a metabolic shift away from oxidative phosphorylation toward glycolysis^54^. We would expect this to be reflected in changes in respiratory chain organization, a hypothesis we could now test *in situ*. We performed the same three-color ExM imaging protocol described above on both non-activated (Fig. 4h,i) and activated (Fig. 4j,k) primary human T-cells. Applying qExM, we found increased isolated complex densities for CI and CIII and decreased CIV density under activated conditions, relative to non-activated. In addition, we measured a significant reduction in super-complex abundance in activated T-cells (Fig. 4l), specifically for Complex I+III+IV configurations, suggestive of a super-complex level regulation of respiratory function during T-cell activation.

## Discussion

The application of qExM to mitochondrial respiratory chain complexes demonstrates its versatility for investigating endogenous targets of unknown abundance. Our measurements of complex densities are the first to generate absolute estimates, while also providing the crucial advantage of spatial context and single organelle information, offering additional nuance to findings from bulk biochemical linvestigations^55^. We also detected and quantified respiratory chain super-complexes in situ, revealing their distribution patterns without disrupting relative positioning. The observation that T-cell activation leads to a significant reduction in the density of fully-assembled super-complexes illustrates how qExM can connect molecular reorganization to functional cellular states.

Previous studies have applied single-molecule localization microscopy (SMLM) to quantify antibody-labeled endogenous targets. While these studies provided valuable insights, they did not take into account imperfect labeling efficiency, leading to underestimated abundances^3,4,56–58^. SMLM has also yielded accurate quantitative measurements of protein abundance for non-endogenous targets, with calibration constructs and engineered labeling systems such as Halo, SNAP, or ALFA tags^1,2,9^. The labeling efficiencies achievable by qExM are comparable to those measured for SMLM (qExM maximum labeling efficiency was 80.5 ± 5.5% for MT-CO1, two-target Exchange-PAINT maximum labeling efficiency was 76.2 ± 8.4% for the combination of labels 1H1 (GFP), 1B2 (GFP), and 1G5 (Alfa)^1^. qExM enables quantitative imaging of endogenous targets with conventional antibody labeling, at a level previously only possible with engineered samples and SMLM. Current limitations of qExM mostly arise from antibody quality, as the accuracy of abundance estimates depends on their specificity. The method also assumes independent binding of the antibodies during dual labeling, so epitopes should be sufficiently distant to avoid interference. The trade-off between expansion factor and antigenicity preservation means that high expansion factors may not be appropriate for quantitative applications, particularly because antibody labeling efficiency is paramount.

The potential applications of qExM extend well beyond the examples presented here. As multiplexed sequential imaging methods continue to advance, this framework could be applied to simultaneously quantify dozens of targets, enabling comprehensive visual super-resolved proteomics^11,59^. Recent extensions of expansion microscopy to whole organisms and diverse tissue types suggest that qExM could be adapted for biological systems unsuited for genetic modification, from developing embryos to clinical pathology samples^23,60^. By providing quantitative information about endogenous protein abundances in situ, qExM creates new opportunities to connect molecular-scale organization with emergent cellular functions.

## Methods

### Computational Simulations

We performed simulations to evaluate the accuracy and precision of two population estimators under varying conditions of labeling efficiency and target abundance. Additionally, the simulations modeled both individual nuclear pore complexes (8 targets per complex) and images containing multiple nuclear pore complexes (320 targets) to provide some indication of the estimators’ performances under different data aggregation strategies.

For each scenario, we generated a population of virtual targets. Each target was independently assigned label status (labeled or unlabeled) for each of two label types by drawing binomially distributed random numbers with probabilities of success corresponding to labeling efficiencies 0.1 or 0.8. We recorded the total number of targets positive for each label and the number of dual-labeled targets.

Estimates for the number of targets were calculated using two different estimators^8^:

1. the Petersen estimator:

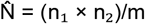
2. the Chapman estimator:

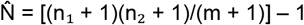

In all formulations, n_1_ and n_2_ represent the number of targets labeled with each probe, and m represents the number of dual-labeled targets.

For each condition we simulated 1,000,000 independent experiments and averaged the results. Performance was assessed using bias, here defined as the expected value of the target count estimates minus the true count.

Bias curves were generated by defining the true number of targets across the range from 5 to 1000 with the labeling efficiency set to either 10% or 80%. For each true count, the Peterson and the Chapman estimators were calculated from 1,000,000 trials and the bias was determined.

### U2OS Cell Culture

U-2 OS cells with CRISPR-edited endogenous NUP96-mEGFP (Cell Line Service, catalog #300174) were cultured in McCoy’s 5A Modified Medium (Gibco) supplemented with 10% fetal bovine serum, 100 U/ml penicillin, and 100 μg/ml streptomycin. Cells were maintained at 37°C in a humidified atmosphere with 5% CO_2_ and used for a maximum of 20 passages.

For imaging experiments, cells were seeded at a density of 50,000 cells per well on uncoated 12 mm #1.5 glass coverslips in 24-well plates. Cells were allowed to adhere and grow for 24-48 hours prior to cryo-fixation to achieve approximately 70% confluence.

### T-Cell Isolation and Activation

Anonymized human buffy coats from three different healthy donors were obtained at the Center of Interregional Blood Transfusion SRK Bern (Bern, Switzerland). Peripheral blood mononuclear cells (PBMCs) were isolated by density gradient centrifugation using Lymphoprep (Accurate Chemical and Scientific Corporation). Non-activated human T cells were isolated using the EasySep™ Human T Cell Isolation Kit (Stemcell techonologies, Cat. # 17951) according to manufactures instructions. Isolated non-activated T cells were either processed immediatedly for qExM or incubated overnight in T Cell medium (TCM) (RPMI 1640 with GlutaMax [Gibco] supplemented with: 10% Fetal bovine serum [cytiva], 10mM HEPES [Gibco], 1mM sodium pyruvate [Thermo scientific], 0.1% 2-mercaptoethanol [merck], 2mM L-glutamine (Thermo Scientific), 1% non-essential amino acids (Gibco), 1% Penicilin streptomycin [PanReac AppliChem] and 100 IU of IL-2 (Peprotech). T-cell activation was performed immediately or the day after T cell isolation in non–tissue culture–treated 24-well plates (Corning) coated with anti-CD3 (OKT3) (BioLegend) and anti-CD28 antibody (BioLegend) in TCM including 100 IU IL-2 for 30h.

### Cryo-fixation, Freeze Substitution, and Expansion Microscopy

#### Cryo-fixation and Freeze Substitution

Cells were prepared for expansion microscopy using a cryo-fixation protocol adapted from Laporte et al^24^. Coverslips with adherent cells were mounted in forceps on a custom-built plunge freezer, and excess liquid was removed by using filter paper. The coverslips were rapidly plunged into a liquid ethane/propane mixture cooled by liquid nitrogen. The frozen coverslips were immediately transferred to 5 mL eppendorf tubes containing 3 mL of dry acetone pre-frozen in liquid nitrogen. These tubes were frozen at approximately 45° angles to maximize surface contact with the coverslips. The sealed tubes were buried in dry ice in a Styrofoam cooler and placed on a rocking shaker for 24 hours.

Next, the tubes were gradually warmed to room temperature over 1.5 hours. The coverslips were then removed from acetone, cell-side identified, and transferred to 12-well plates for sequential rehydration. Samples underwent a graded ethanol series consisting of two 5-minute incubations each in 100% and 95% ethanol, followed by single 5-minute incubations in 75%, 50%, and 25% ethanol, and finally three washes in PBS to completely remove residual ethanol.

In the sample preparation comparison, the samples were fixed for 15 minutes at room temperature in PBS with 3% formaldehyde and 0.1% glutaraldehyde.

#### Sample Anchoring

Prior to expansion, cellular proteins gained the anchoring functional group using glycidyl methacrylate (GMA). Samples were washed three times with 100 mM sodium bicarbonate buffer and then incubated with 0.04% (v/v) GMA in 100 mM sodium bicarbonate for 4 hours at room temperature on a rocking shaker. Following anchoring, samples were washed three times with PBS to remove unbound GMA.

In the sample preparation comparison, formaldehyde-acrylamide anchoring was conducted by incubating the samples in 1.4% formaldehyde, 2% acrylamide in PBS at 37^°^ for 5 hours.

#### Gel Embedding and Expansion

The expansion monomer solution was prepared with 23% (w/v) sodium acrylate, 10% (w/v) acrylamide, and 0.1% (w/v) N,N’-methylenebisacrylamide (BIS). The sodium acrylate was prepared as a 46% solution from powder stored under inert gas. The complete monomer solution was degassed under vacuum for 1 hour before use. Coverslips with anchored cells were pre-incubated in the monomer solution without initiators to ensure complete equilibration for 20 minutes at room temperature on a rocking shaker.

For gelation, a casting chamber was prepared using a glass slide in a petri dish pre-chilled at −20°C. Working on ice, tetramethylethylenediamine (TEMED) and ammonium persulfate (APS) were added to the monomer solution to final concentrations of 0.5% each. Coverslips with monomer solution were placed in a humidified chamber and polymerized at 37°C for 1 hour.

Following polymerization, the gels with attached coverslips were immersed in denaturation solution (200 mM SDS, 200 mM NaCl, 50 mM Tris, pH 9.0) at 95°C in 12-well plates. After approximately 5 minutes when the gels detached from the coverslips, they were transferred to 1.5 mL eppendorf tubes with fresh 95°C denaturation solution and incubated at 95°C for an additional 1 hour 45 minutes. The gels were then placed in 250 mL of deionized water to remove SDS and initiate expansion.

#### Immunostaining of Expanded Samples

The expanded gels were sectioned using the cap of a 5 mL eppendorf tube to create manageable subsections, which were transferred to PBS. For immunostaining, gel subsections were incubated in 100 μL of blocking buffer (3% BSA in PBS) containing primary antibodies at 1:100 dilution in 2 mL eppendorf tubes. The samples were incubated overnight at 30°C with gentle agitation.

Following primary antibody labeling, samples were washed three times with 5-minute incubations in blocking buffer. Secondary antibodies were then applied at 5 μg/mL in blocking buffer and incubated overnight at 30°C with agitation. After immunolabeling, samples were washed twice in PBS and re-expanded in 50 mL of deionized water for 30 minutes.

To stabilize the antibody labeling, samples were post-fixed with 6% formaldehyde in water for 30 minutes, followed by extensive washing in deionized water. The cell-containing side of each gel was identified by examining the sample on the microscope and mounted on poly-D-lysine (PDL) coated coverslips. These coverslips were prepared by sequential sonication for five minutes in 1M KOH and ethanol, followed by incubation with PDL solution for at least 1 hour at 37°C, and a final rinse in water to remove excess PDL.

#### Antibody Conjugation

Custom fluorophore-conjugated secondary antibodies were prepared for two- and three-color STED microscopy experiments. Secondary antibodies from Jackson ImmunoResearch (100 μL containing 120-130 μg protein) were buffer-exchanged into 100 mM sodium bicarbonate using Zeba spin desalting columns (40K MWCO, Thermo Fisher Scientific). The antibodies were then reacted with a 15-fold molar excess of NHS-ester fluorophores (1 μL of 10 mM stock solution of either Alexa Fluor 488, Atto490LS, Alexa Fluor 594, or Star635P in anhydrous DMSO).

Conjugation reactions proceeded for 4 hours at room temperature with gentle agitation in the dark. Unreacted fluorophores were removed by a second buffer exchange using fresh Zeba spin desalting columns, equilibrated with PBS containing 0.1% sodium azide as a preservative. The final antibody concentration and degree of labeling (DOL) were determined spectrophotometrically using a NanoDrop spectrophotometer. All conjugated antibodies used in this study had DOL values between 2 and 4 fluorophore molecules per antibody, ensuring fluorescence while minimizing potential detriment to the binding affinity of the antibody.

#### Primary antibodies

In this study we used the following primary antibodies:

for CI: NDUFS2 (Abcam ab192022), ND2 (Sigma MABS2047), for CIII: UQCRCP1 (Abcam ab14746), UQCRFS1/RISP (Abcam ab11925), for CIV: MT-CO1 (Abcam ab14705), MT-CO2 (Abcam ab79393), GFP (Abcam ab6673), Nup96 (ProteinTech-12329-1-AP), Nup54 (Sigma HPA035929), ELYS (Sigma HPA031658), P5CS (Proteintech 17719-1-AP, 68184-1-IG)

All antibodies were used at a dilution of 1:100, except ELYS which was used at a concentration of 1:50.

#### STED Microscopy and deconvolution

Expanded samples immobilized on poly-D-lysine coated coverslips were imaged using a Leica SP8 STED microscope equipped with a HC PL APO 93×/1.30 NA glycerol immersion objective with a motorized collar correction ring for spherical aberration compensation. Images were acquired with a pixel size of 13.5 nm in either 1848 × 1848 or 1152 × 1152 pixel fields of view. All images were collected with a pixel dwell time of 2.15 μs, with 8 line accumulations.

For multicolor STED imaging, sequential scanning by line was employed to minimize channel crosstalk. The depletion laser (775 nm) power was optimized for each fluorophore to achieve maximum resolution while minimizing photobleaching, and kept consistent across all images. Excitation of Star635p was done with 633 nm illumination, Alexa Fluor 594 with 561 nm illumination, and Atto490LS and Alexa Fluor 488 with 488 nm illumination. Star 635p was detected in the range of 643-750 nm, Alexa Fluor 594 in the range of 580-635 nm, and Atto490LS in the range 580-750 nm.

Raw STED images were deconvolved using Huygens Professional software (Scientific Volume Imaging) with deconvolution parameters including, the use of a theoretical PSF, STED saturation factor of 25, STED immunity of 5, classic maximum likelihood estimation as the algorithm, signal/noise of 15, legacy option for acuity, automatic background estimation, stopping criteria of 1 iteration with a 0.01 quality change.

#### NPC model depiction

NPC atomic model depiction was generated using PDB-7R5J in ChimeraX.

#### Nuclear pore identification

Nuclear pores were identified manually using a systematic visual inspection approach. Deconvolved multicolor STED images were imported into napari, the open-source multi-dimensional image viewer^61^. The center coordinates of each identifiable nuclear pore complex were marked by an experienced observer and recorded. These coordinates were exported as CSV files for each image to facilitate subsequent analysis.

To ensure consistency in nuclear pore identification, we applied the following criteria: nuclear pores had to display the characteristic ring-like or punctate pattern with signal intensity significantly above the local background. Incomplete or ambiguous structures were excluded from analysis to minimize false positives.

#### Mitochondria segmentation

Mitochondrial structures were isolated from the multicolor images using a semi-automated segmentation approach. First, we generated composite images by summing the signal intensities across the separate fluorescent channels to maximize mitochondrial signal representation. These composite images were then processed with a Gaussian blur filter to reduce noise while preserving structural features.

The preprocessed images were segmented using a machine learning classifier trained in Labkit ^42^ (ImageJ plugin). The classifier was trained on a set of manually annotated mitochondrial structures. The trained classifier was then applied to the full dataset to generate binary segmentation masks. These final segmentation masks were stored as TIFF files with each mitochondrion represented as a distinct labeled region. Quality control was performed by visual inspection of the segmentation results overlaid on the original images.

#### Image Instance Segmentation and Puncta Analysis

Deconvolved STED images were processed through an automated instance segmentation pipeline using CellProfiler^36^ (version 4.2.7) to identify and characterize individual fluorescent puncta representing labeled proteins. The processing workflow consisted of the following sequential steps:

1. **Intensity Normalization** Pixel intensity values were rescaled to a standardized range (0-1) to ensure consistent thresholding across different images and acquisition sessions.
2. **Image Smoothing** Images were smoothed with a gaussian filter and artifact diameter of 2 pixels.
3. **Primary Object Identification** Puncta were identified as primary objects with diameter constraints between 1 and 50 pixels with a global threshold strategy.
4. **Clumped Object Separation** Closely spaced puncta were distinguished by intensity-based watershed segmentation.
5. **Feature Extraction** For each identified object, we extracted morphological and intensity features including centroid coordinates, integrated intensity, and area.

The segmentation results were exported as CSV files containing the x-y coordinates of each objects’s centroid along with associated feature measurements. These centroid coordinates were subsequently used for spatial analysis of protein distribution patterns and colocalization studies. The complete CellProfiler pipeline is available on the Github page.

#### Colocalization Threshold Determination

Colocalization distance thresholds between fluorescent signals from different channels was determined using a data-driven approach nearly identical to that which was previously introduced for SMLM^1,31^. For NPC analysis, relevant puncta were defined as those within a 25-pixel (~82 nm) radius from the manually identified NPC centers. For mitochondrial targets, relevant puncta were constrained to those located within the segmented mitochondrial masks.

For each image, we calculated the first nearest neighbor distance (1-NND) between every object in one channel and all objects in the second channel using k-dimensional tree (KD-tree) spatial indexing for computational efficiency. For three color data, this process was done between each pair of channels.

Next, we performed five independent simulations where one channel’s objects were kept in their original positions while the second channel’s objects were randomly distributed within the same spatial constraints (NPC center vicinity or mitochondrial mask).

Following ^1,31^, we varies he mean and standard deviation of the randomly applied offsets as well as the fraction of colocalized objects. The best fit parameter values minimized the sum of the squared difference between the experimental and simulated 1-NND distributions. The final colocalization cutoff distance was defined as the mean offset distance plus one standard deviation.

**Table 1.** Simulation defined cutoff distance – Distance in expanded space.

| Label Pair | Mean Offset (nm) | Offset Standard Deviation (nm) | Cutoff distance (nm) | Cutoff (divided by expansion factor) (nm) |
| --- | --- | --- | --- | --- |
| NPC<br>mEGFP –<br>nup96 | 74.69 | 33.95 | 108.64 | 25.26 |
| NPC<br>mEGFP –<br>nup54 | 101.1 | 29.48 | 130.57 | 30.36 |
| NPC<br>mEGFP –<br>ELYS | 103.49 | 32.09 | 135.58 | 31.53 |
| Mitochondria<br>CI | 121.99 | 51.47 | 173.45 | 40.33 |
| Mitochondria<br>CIII | 103.07 | 47.17 | 150.24 | 34.94 |
| Mitochondria<br>CIV | 104.03 | 48.2 | 152.23 | 35.40 |
| Mitochondria<br>CI-CIII | 74.42 | 55.69 | 130.11 | 30.26 |
| Mitochondria<br>CI-CIV | 103.77 | 51.78 | 155.55 | 36.17 |
| Mitochondria<br>CIII-CIV | 91.45 | 48.47 | 139.93 | 32.54 |

This cutoff was then applied back to the original data, where object pairs with inter-channel distances below the threshold were classified as colocalized.

### Quantitative Analysis of Labeling Efficiency and Target Abundance

For each image, we quantified the key variables needed for abundance estimation: the total number of detected objects in each channel (n_1_, n_2_) and the number of colocalized object pairs (m) identified using the distance-based thresholds established previously.

Labeling efficiencies were calculated as:

Labeling efficiency of probe 1 (E_1_) = m/n_2_

Labeling efficiency of probe 2 (E_2_) = m/n_1_

The total number of target proteins was estimated using the Chapman estimator, which reduces bias in scenarios with small sample sizes or low colocalization rates.

See Methods: Computational Simulations and Fig. S1 for more details.

#### Measurement of P5CS containing fibers

U2OS nup96-mGFP cells were subjected to 8 hours of galactose media consisting of DMEM, no glucose (Thermo Scientific 11966025), Dialyzed FBS (Thermo Scientific A3382001), 10 mM galactose (powder), Pen/Strep, 0.1 mM NEAA (Thermo Scientific 11-140-076), 50 ug/ml uridine, 2 mM L-glutamine, 1 mM Sodium Pyruvate (Thermo Scientific 11360070). Samples then were cryofixed and expanded as discussed previously. For complex I and complex IV experiments, P5CS filaments were labeled with P5CS antibody (Proteintech 68184-1-IG), and for complex III experiments P5CS filaments were labeled with P5CS antibody (Proteintech 17719-1-AP). In addition to the secondaries to label the appropriate complex target, these samples were co-labeled with secondary antibody conjugated to Alexa Flour 488 to identify the P5CS domain.

To identify mitochondrion in the data with and without the P5CS fiber, mitochondria were hand segmented into these two groups through the use of napari^61^ to define the different regions.

An example image was captured labeled for P5CS and Mt-CO1 on a Nikon NSPARC in SR mode, Galvano scanner, 2x line integration, 0.5 µsec dwell time, Plan Apo IR 60x WI DIC N2 objective, excitation 561 nm and 488 nm and emission range 570-616 nm and 502-546 nm. Imaged was deconvolved over 20 iterations with Richardson-Lucy deconvolution in DeconvolutionLab2^62^.

#### Determination of Nuclear Pore Complex Protein Labeling Efficiency

We employed a maximum likelihood estimation approach to quantify the effective labeling efficiency of individual proteins within nuclear pore complexes (NPCs) as previously described^30^. This established approach accounts for the eight-fold symmetry of NPCs, where each subunit contains a defined number of binding sites for the target proteins.

For each NPC, we recorded the number of subunits (0-8) that contained at least one labeled protein. These observed distributions were analyzed across three nucleoporin datasets: Nup96-GFP imaged with Nup54, Nup96, or ELYS. The probability of visualizing a subunit depends both on the protein-level labeling efficiency and the number of target proteins present in each subunit.

For each nucleoporin dataset, we minimized the negative log-likelihood for observing a number of labeled subunits as a function of the protein-level labeling efficiency. The minimization was implemented in Python using the minimize_scalar function from the SciPy package. The value of the “method” parameter was “bounded.”

#### Correcting for Incomplete Labeling in Respiratory Chain Super-Complex Analysis

To account for incomplete labeling in our three-color expansion STED images, we used a linear algebra-based approach that enables quantification of respiratory chain super-complex abundance. This method specifically addresses the systematic undercounting that occurs when target proteins escape detection due to incomplete labeling efficiency.

For our three-channel imaging system (CI, CIII, and CIV), we defined seven possible configurations for protein complexes:

1. Isolated CI
2. Isolated CIII
3. Isolated CIV
4. CI-CIII pairs
5. CI-CIV pairs
6. CIII-CIV pairs
7. CI-CIII-CIV triplets

The relationship between observed counts (O) and the estimated abundance (N) was modeled using a linear system:

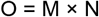

where M is a 7×7 matrix encoding the probabilities of observing each possible labeled state given a true underlying configuration. It follows that N is a 7 x 1 vector containing the estimators for the true abundance of each configuration defined above.

Each element M[i,j] represents the probability that a complex in true state j will be observed as having labeling outcome i due to some labels failing to attach. These probabilities were calculated using the experimentally determined labeling efficiencies (LE) for each channel:

- For example, the probability that a true CI-CIII-CIV triplet appears as an isolated CI signal is: M[CI, CI-CIII-CIV] = (LE_CI) × (1 - LE_CIII) × (1 - LE_CIV)
- Similarly, the probability of observing a CI-CIII pair when the true state is a CI-CIII-CIV triplet is: M[CI-CIII, CI-CIII-CIV] = (LE_CI) × (LE_CIII) × (1 - LE_CIV)

The complete matrix incorporated all possible observation scenarios based on the combinatorial outcomes of successful and unsuccessful labeling events. Notably, it is upper triangular because we assume that the probability of measuring a label where there was no matching target complex was zero. This and the fact that its values are bound to the range [0, 1] with labeling efficiencies greater than 10% help ensure that it is invertible and well-conditioned.

The true counts for each configuration were reconstructed from the observed counts by solving the linear system:

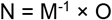

where M^-1^ is the inverse of the matrix M. To account for variations in mitochondrial content across samples, the absolute abundance values were normalized to the mitochondrial area by dividing the number of objects by the area of the corresponding mitochondrial mask (in μm^2^). This yielded density measurements (complexes/μm^2^) that could be compared across different samples and experimental conditions.

### Expansion factor size determination

#### Nuclear Pore Complex as a Dimensional Reference Standard

We utilized the well-characterized dimensions of nuclear pore complexes (NPCs) to determine the expansion factor achieved in our expansion microscopy protocol. Specifically, we used Nup96-mEGFP labeled NPCs as intrinsic dimensional standards, since their native dimensions have been precisely determined by cryo-electron microscopy studies.

#### Measurement Methodology

To obtain accurate measurements of expanded NPCs, we implemented the following systematic approach:

1. We identified completely labeled NPCs where all eight subunits were identified by mEGFP, ensuring the most accurate dimensional assessment.
2. For each selected NPC, we measured the maximum diameter by determining the longest distance between opposing subunits. This approach minimized measurement errors that could arise from the 2D projection of NPCs positioned along curved nuclear membranes, which can appear as elliptical rather than circular structures.
3. Multiple measurements were taken across different NPCs and from multiple cell samples to establish a statistically robust estimate of the expanded NPC diameter.

#### Calculation of Expansion Factor

The expansion factor (EF) was calculated as:

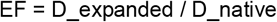

Where:

- D_expanded is the measured diameter of NPCs after expansion (in nm)
- D_native is the established diameter of NPCs from cryo-EM literature (104 nm for the Nup96 scaffold ring)

This approach provided a precise, sample-specific calibration of the expansion factor that could be applied to all subsequent measurements within the same expanded sample. The calculated expansion factor had a mean of 4.3x.

#### Signal to Noise Measurements

To assess the difference in antibody labeling across the experimental condition, signal to noise measurements were obtained by measuring the difference in signal between the maximum signal from the brightest MT-CO1 labeled mitochondria and the unlabeled background and dividing this value by the standard deviation of the background.

#### False Positive Measurements

To compare false positive labeling, U2OS nup96-mGFP cells were cryo fixed and subjected to freeze substitution as before. The samples were blocked in a blocking buffer composed of 3% BSA in PBS for 1 hour, stained with primary antibody for GFP (1:500), washed 3x with 5-minute incubation in blocking buffer, stained with secondary antibody (2 µg/mL) for donkey anti goat conjugated to Alexa Flour 488 and imaged with a Nikon Spinning Disk confocal W1. The objective used was SR HP Apo TIRF 100x 1.49 NA oil objective. Illumination was 488 nm, pixel size 110 nm, 100 ms exposure time.

#### Larger gels

Larger gels were formed by using a monomer solution containing 28% SA, 10% AA, 0.05% bis, 4% DMAA, and denatured in 10% w/v SDS, 8M Urea, 25 mM EDTA, 2x PBS, pH 7.5 for 72 hours at 80°C shaking^33^. Gels were stained as discussed earlier for Mt-CO1, with secondary antibodies conjugated to CF568, and imaged with a Nikon Spinning Disk confocal W1. The objective used was a PLAN APO IF 60x 1.27 water immersion objective.

## Code Availability

Data and code used in this study are deposited here: https://github.com/LEB-EPFL/qExM_code

## Acknowledgements

We thank Hélène Perreten for the U2OS-mEGFP cell culture work, Mark Bates for the discussions on quantitative imaging and antibody recommendations for the NPC, Christiane Beninca for antibody recommendations for mitochondrial proteins, and Max Kieff for discussions on statistics. This work was supported by the Swiss National Science Foundation project grant (SNSF, 310030_215737, S.M.), the Human Frontiers in Science Program (HFSP, RGP0038/2021, S.M.), and the European Union’s H2020 program under the European Research Council (ERC, CoG 819823 Piko, S.M.).

